# A Small-Molecule Mitochondrial Complex I Modulator Improves Behavioral and Mitochondrial Dysfunction in Schizophrenia

**DOI:** 10.64898/2026.05.19.726440

**Authors:** Maltesh Kambali, Sergey Trushin, Muxiao Wang, Rajasekar Nagarajan, Jinrui Lyu, Eugenia Trushina, Uwe Rudolph

## Abstract

Weak inhibition of mitochondrial complex I (mtCI) has been shown to have neuroprotective effects in cellular and animal models of Alzheimer’s and Huntington’s diseases, at least in part by enhancing mitochondrial biogenesis and function. Mitochondrial dysfunction has also been demonstrated in schizophrenia patients and mouse models of schizophrenia. We tested whether weak inhibition of mtCI would ameliorate mitochondrial and behavioral phenotypes in a mouse model of schizophrenia. In mice with four copies of the *Gldc* gene, 8 weeks of treatment with the weak mtCI inhibitor, the small-molecule tricyclic pyrone compound CP2, reversed spontaneous alternation deficits in the Y maze, startle habituation deficits, and social novelty deficits in the three-chamber social interaction test. Consistent with the mechanism of action, Western blots revealed that CP2 reverses the reduced expression of PGC-1α, a master regulator of mitochondrial biogenesis, and of the VDAC1, a primary gatekeeper for the exchange of metabolites, ions, and ATP between mitochondria and the cytosol. These findings suggest that the improvement of mitochondrial function may represent a novel strategy to reverse pathophysiological and behavioral deficits in schizophrenia.

## Introduction

Schizophrenia is a complex neuropsychiatric disorder whose symptoms may include cognitive impairment, disrupted sensory processing, and social dysfunction. While traditionally associated with dopaminergic and glutamatergic dysregulation^1,2^, converging evidence over the past two decades has implicated mitochondrial dysfunction as a central contributor to disease pathophysiology^3–5^. This shift reflects a broader view of schizophrenia as a disorder of cellular energy metabolism and neural circuit stability rather than solely neurotransmitter imbalance.

Postmortem and clinical studies in patients with schizophrenia consistently report abnormalities in mitochondrial structure and function^6^. Reductions in mitochondrial density, altered cristae morphology, and impaired oxidative phosphorylation have been reported in brain regions critical for cognition, including the prefrontal cortex and hippocampus^7,8^. In parallel, gene expression studies, transcriptomic analyses, and multi-omic analyses reveal downregulation of genes involved in electron transport chain activity, ATP production, and mitochondrial biogenesis, including reduced expression of regulators such as PGC-1α. Functional imaging studies further support these findings, showing altered cerebral energy metabolism and decreased bioenergetic efficiency in individuals with schizophrenia^9,10^. Together, these data suggest that impaired mitochondrial function contributes to synaptic deficits, reduced plasticity, and network-level dysfunction underlying cognitive and behavioral symptoms.

Animal models have provided mechanistic insights into how mitochondrial dysfunction may drive schizophrenia-relevant phenotypes. Rodent models with genetic or pharmacological disruption of mitochondrial pathways exhibit deficits in working memory, sensorimotor gating, and social interaction, mirroring core features of the disorder^11–15^. For example, impairments in mitochondrial oxidative phosphorylation have been linked to altered glutamatergic and dopamine signaling^16,17^. Similarly, models with disrupted mitochondrial biogenesis or increased oxidative stress show abnormalities in synaptic transmission and neuronal connectivity^6^. Importantly, these models also demonstrate that restoring mitochondrial function can rescue behavioral deficits, supporting a causal role for bioenergetic dysfunction.

Therapeutic strategies targeting mitochondrial pathways have therefore gained increasing attention^18–21^. Interventions aimed at enhancing mitochondrial biogenesis, improving oxidative phosphorylation, or reducing oxidative stress have shown promise in both preclinical and clinical settings. Compounds that activate AMP-activated protein kinase (AMPK), increase NAD^+^ availability, or upregulate PGC-1α signaling have been associated with improved cognitive function and synaptic plasticity in models of neuropsychiatric and neurodegenerative diseases^19–21^. However, many of these approaches have not been evaluated in animal models for schizophrenia. Novel targets and translationally viable strategies are urgently needed.

We have previously demonstrated that brain penetrable small molecule tricyclic pyrone compound CP2 activates adaptive stress response program via weak inhibition of mtCI^20,21^. By modestly reducing complex I activity, CP2 increases the AMP/ATP ratio, leading to activation of AMPK and downstream pathways that enhance mitochondrial biogenesis, autophagy, and cellular stress resistance^20,21^. This mechanism is consistent with “mitohormesis,” in which controlled metabolic stress promotes long-term cellular resilience. Preclinical studies have demonstrated that CP2 improves mitochondrial function, synaptic activity, and cognitive performance across multiple disease models. In models of Alzheimer’s disease, chronic CP2 treatment restored long-term potentiation, reduced oxidative stress, blocked the ongoing neurodegeneration and improved memory performance^20^. Chronic CP2 administration to mice was safe over 14 months^22,23^. Similarly, in Huntington’s disease models, CP2 enhanced neuronal survival and normalized energy metabolism^24,25^. These studies highlight CP2’s ability to broadly modulate mitochondrial and neuronal function *in vivo*, with sustained behavioral benefits following chronic administration.

Despite these advances, the potential of CP2 in neurodevelopmental and schizophrenia-relevant models has not been fully explored. Given the strong evidence linking mitochondrial dysfunction to schizophrenia-related phenotypes, we hypothesized that CP2-mediated metabolic reprogramming could ameliorate behavioral and molecular deficits in the recently developed mouse model with 4 copies of the *Gldc* gene (hereafter called 4cG mouse model)^12,13^. In this model, an increased copy number of the *Gldc* gene encoding the glycine-degrading enzyme glycine decarboxylase results in reduced extracellular concentrations of the NMDA receptor co-agonist glycine in the dentate gyrus. The 4cG model, characterized by impairments in defined biochemical pathways, mitochondrial oxygen consumption rate, long-term potentiation, startle habituation, social interaction behavior, and cognition, provides a relevant system in which mitochondrial dysfunction is likely to play a key role. Deficits in startle habituation, spontaneous alternation, and sociability can be reversed in this model with chronic administration of glycine^26^. Similarly, acute administration of lactate reversed the deficits in startle habituation, spontaneous alternation, and social preference in the 4cG mice, as well as the reduced expression of PGC-1α and BDNF^27^.

Based on CP2’s established mechanism of action and its demonstrated efficacy in restoring mitochondrial and synaptic function, we investigated whether chronic CP2 treatment could reverse deficits in mitochondrial biogenesis mechanisms, working memory, sensory processing, and social interaction behavior in 4cG mice. This approach allowed us to directly test the hypothesis that targeting mitochondrial bioenergetics is sufficient to rescue complex behavioral phenotypes associated with schizophrenia-relevant pathology.

## Methods and Materials

### Animals

Mice with four copies of glycine decarboxylase (4cG) and wild type control mice on the C57BL/6J background mice were bred and maintained in standard conditions (temperature 23°C, 12/12-h light/dark cycles with lights on at 7:00 am) with free access to food and water. All experimental procedures described in this paper were in strict accordance with the National Research Council’s Guide for the Care and Use of Laboratory Animals and were approved by the Institutional Animal Care and Use Committee (IACUC) of University of Illinois, Urbana-Champaign. Adult males and females (2-4 months of age) were used in this study.

### CP2 treatment

CP2 was synthesized by Nanosyn, Inc. (http://www.nanosyn.com), as described previously,^28^ and was purified using HPLC. Authentication was performed through NMR spectra to ensure the lack of batch-to-batch variation in purity. CP2 was synthesized as a free base. Mice received CP2 (25 mg/kg) or vehicle (PEG400) either as a single oral gavage dose for acute studies or in drinking water *ad lib* for 8 weeks for chronic behavioral and molecular studies.

### Y-maze spontaneous alternation

Working memory was measured using a symmetrical Y-maze comprising three arms (30 cm × 8 cm × 15 cm) oriented at 120° relative to one another. The experiment was performed using a previously published protocol with minor modifications^29–31^. Mice were introduced at the end of one arm (Arm A or Arm B) and permitted to explore freely for 8 minutes. The sequence (pattern) and total number of arms entered were recorded. An arm entry is considered to be complete when the hind paws of the mouse had been completely placed in the arm. The percentage of alternations was calculated as the number of alternations divided by the total number of eligible arm-entry triads-2, multiplied by 100. As the re-entry into the same arm was not counted for this analysis, the chance performance level in this task was 50% for the choice between the arm mice visited more recently (non-alternation) and the other arm the mice visited less recently (alternation).

### Prepulse inhibition and startle habituation

Prepulse inhibition (PPI) was determined using the SM100 Startle Monitor System (Kinder Scientific, USA), consisting of a sound-attenuated startle chamber and Startle Monitor software. Animals were placed in an adjustable Plexiglas holder, allowing limited movement but not providing restraint, positioned directly above a sensing platform. Each animal underwent 15 min of habituation with the holder for two days in the home cage, followed by habituation to the testing chamber with white noise for 5 min on the next day. On test day, each test session consisted of 5 min acclimatization with background white noise (70 dB), followed by presentation with 6 startle pulses of 120 dB [intertrial interval (ITI): 20 s]. Animals were then subjected to five types of trials presented 12 times each in a pseudorandom manner: pulse alone, prepulse + pulse, or prepulse alone (prepulse 3, 6, 9 or 12 dB above background). ITI varied from 10 to 20 s (average 15), two stimulus onset asynchrony (SOA) intervals were 60ms and 120 ms, prepulse length 20 ms, and pulse length 40 ms. Each PPI session ended with 6 startle pulse presentation of 120 dB. The PPI test lasted about 35 min. PPI was calculated as % PPI for each prepulse intensity as: [(pulse_alone - prepulse + pulse/pulse alone) × 100]; with a lower percentage indicating decreased PPI. Startle Habituation was assessed by comparing the averages of first (6) and last (6) sets of startle pulse alone trials (first and last response).

### Three-Chamber Social Interaction Test

Sociability and social novelty were evaluated in a three-chamber social interaction test. The apparatus is a rectangular three-chamber box in which each chamber is 20 × 45 cm with a height of 40 cm. Walls and dividing walls are made from clear Plexiglas, with a sliding door middle section with 5 cm width and 5 cm height which allows free access to each chamber. The experiment was performed using a previously published protocol with minor modifications^13,27,32^. The subject mice were habituated to the central compartment with closed doors for 5 min. After the habituation phase, subjects were tested in the sociability task that consisted of exploring either a conspecific stranger mouse 1 (Sex-matched one-month younger mouse inside an inverted cup) or an empty inverted cup located in the opposite compartment with an exploration time of 10 min. After this session and a 5 min break, the social novelty task was performed for 10 min. The empty inverted cup is replaced with an inverted cup holding a novel stranger mouse 2. Mice under the cups are sex-matched to and approximately one month younger than the experimental mice. The apparatus is cleaned with 70% ethanol after each trial. The time spent in each compartment was measured. The data were recorded and analyzed using the EthoVision XT15 (Noldus Inc., Leesburg, VA, USA).

### Western Blots

Hippocampal tissue samples were minced and homogenized in RIPA lysis buffer (Thermo Fisher Scientific, cat no. 89900) supplemented with protease and phosphatase inhibitor cocktails. Homogenates were centrifuged at 15,000 × g for 10 min at 4 °C, and the resulting supernatant was collected as the soluble lysate samples. Protein concentrations were subsequently determined using a BCA protein assay kit. The experiment was performed using a previously published protocol with minor modifications ^27,30,31^. Western blot analyses were performed with n=4 per genotype. Prior to gel loading, lyaste samples were heated to 95°C for 5 minutes, 30 ug of lysate protein samples (along with protein ladder in first well) were electrophoretically separated on an SDS 10-11% polyacrylamide gel and transferred to a precisely cut PVDF membranes. PVDF membranes (Immobilin-P, Millipore Inc., St. Louis, MO, USA) were blocked with 5% skimmed milk powder (LP0031, OXOID, KS, USA) in 0.05% Tween-20/Tris-buffered saline and then incubated with primary antibody overnight in a cold room (∼4°C). After incubation with goat anti-rabbit (1:5000; Cell Signaling) or rabbit anti-mouse (1:3000; Cell Signaling) horseradish peroxidase-conjugated secondary antibodies, for 1 hour and washed with TBS. Semi-quantitative assessment of protein bands was performed by developing the blots for 2 min with chemiluminescence substrate using SuperSignal™ West Femto Maximum Sensitivity Substrate (Thermo Fisher Scientific, Waltham, MA, USA) and imaged the blots that was executed by Protein Simple Fluor Chem R (Protein Simple, USA). Densitometric quantification was performed using Image-J software. Each blot was used to detect multiple proteins. The following primary antibodies were used:

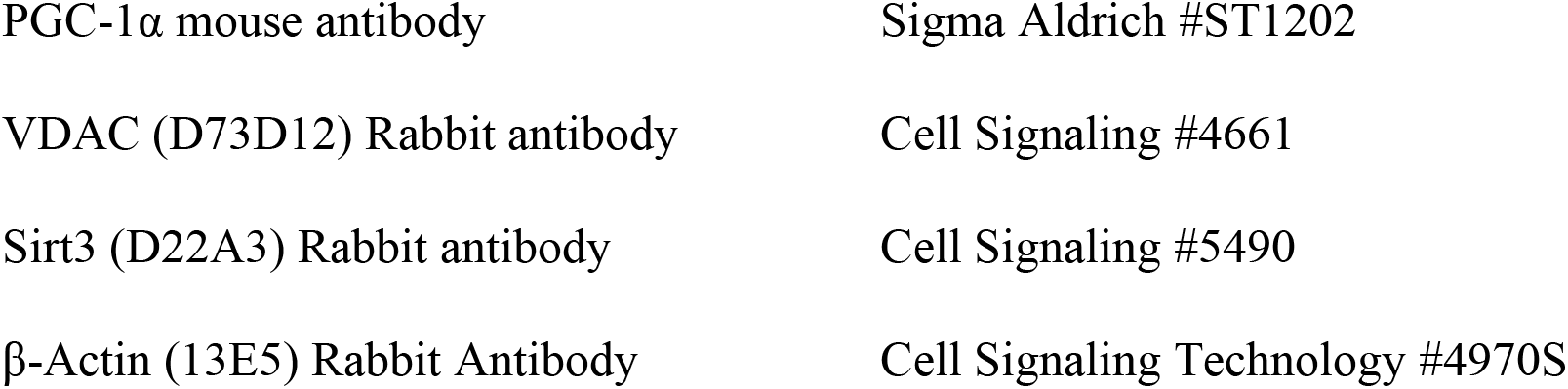

### Statistical Analysis

Sample sizes were based on established practice in respective assays and, to some extent, limited by the availability of mice. Males (2-4) and female mice (3-5) were used in this study. For behavioral tests and Western blots, the statistical analysis was performed using Prism 9 (GraphPad Software, USA). Results were analyzed using one-way ANOVA with post-hoc test Bonferroni’s multiple comparisons method or two-way repeated-measures ANOVA, followed by multiple comparisons with Bonferroni’s correction where appropriate (i.e., only when a significant [p < 0.05] main effect was detected). The significance alpha value is set at < 0.05 for all tests.

## Results

In mouse models of Alzheimer’s disease, behavioral effects of CP2 have been observed from 2-3 months of treatment^20^. To determine the time course of treatment in the 4cG model for schizophrenia, we first evaluated the effects of a CP2 dose of 25 mg/kg body weight on working memory in the Y-maze spontaneous alternation test. Behavioral assessments were conducted in two-week intervals across the treatment course **(Fig. 1A)**. 4cG mice exhibited reduced spontaneous alternation performance compared to wild-type (WT) controls at baseline, consistent with working memory deficits **(Fig. 1B)**. The CP2 treatment group also showed significantly reduced percent alternations at 2 weeks evaluation (**Fig. 1B**). At 4 weeks after treatment, CP2-treated 4cG mice performed similar to the WT mice (**Fig. 1D**). Chronic CP2 treatment led to a progressive improvement in performance, with a significant restoration of spontaneous alternation after 8 weeks of treatment in 4cG mice compared to vehicle-treated 4cG mice **(Fig. 1H)**. At all-time points, the animals showed no change in the total number of entries into the arms of Y-maze (**Fig. 1C, E, G, I)**. Since significant differences between 4cG mice treated with vehicle or CP2 were observed after 8 weeks of treatment, this time was selected for all subsequent experiments.

**Figure 1.**
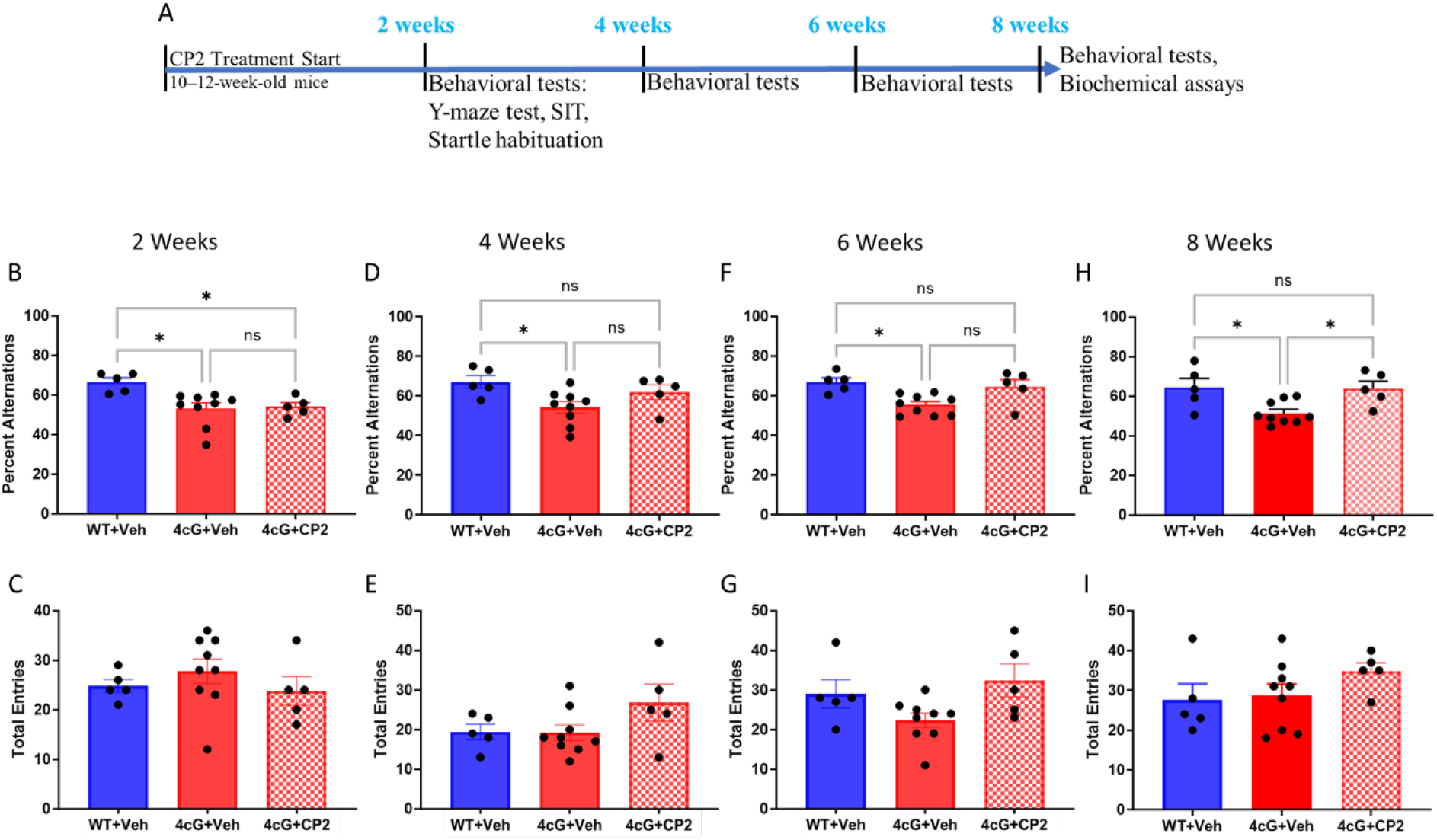
CP2 effects in the Y-maze spontaneous alternation test as a function of CP2 treatment duration in 4cG mice. **A.** Working memory performance was compared between vehicle-treated and CP2-treated groups. **B, C**. Two weeks of CP2 treatment in 4cG mice. The 4cG mice displayed reduced alternations compared to wild type (WT) mice. A one-way ANOVA (F(2,16): 4.515, p=0.0279) followed by Bonferroni’s multiple comparisons test revealed a significant difference between vehicle-treated 4cG mice and vehicle-treated WT mice (t= 2.898, p= 0.0311) and between CP2-treated 4cG mice and vehicle-treated WT mice (t= 2.807, p= 0.0375)]. **D, E**. After four weeks of CP2 treatment, 4cG mice were no longer different from the WT mice. A one-way ANOVA (F(2,16): 4.304, p=0.0319) followed by Bonferroni’s multiple comparisons test revealed a significant difference between vehicle-treated 4cG mice and vehicle-treated WT mice (t= 2.838, p= 0.0352). **F, G** Six weeks of CP2 treatment in 4cG mice. A one-way ANOVA (F(2,16): 6.799, p=0.073) followed by Bonferroni’s multiple comparisons test revealed a significant difference between vehicle-treated 4R mice and vehicle-treated WT mice (t= 3.342, p= 0.0124). **H, I**. Eight weeks of CP2 treatment increased the percent alternations in 4cG mice compared to vehicle-treated 4cG mice. A one-way ANOVA (F(2,16): 5.435, p=0.0158) followed by Bonferroni’s multiple comparisons test revealed a significant difference between vehicle-treated 4cG mice and vehicle-treated WT mice (t= 2.867, p= 0.0335) and a significant difference between vehicle-treated 4cG mice and CP2-treated 4cG mice (t=2.562, P=0.0459). WT+Veh: n=5; 4cG+Veh: n=9, 4cG+ CP2; n=5. All data presented as mean ± SEM.

Next, we assessed whether chronic CP2 treatment rescues schizophrenia-related deficits in information processing and behavioral plasticity, specifically startle habituation to repeated acoustic stimuli and prepulse inhibition of acoustic startle. Vehicle-treated 4cG mice had no prepulse inhibition deficits compared to WT mice at two different stimulus onset asynchronies (SOAs), i.e., 60 ms and 120 ms **(Fig. 2A, B)**, and there were no changes with the CP2 treatment **(Fig. 2A, B)**. Therefore, we analyzed startle habituation, i.e., the responses to the startle pulses presented before and after the prepulse inhibition protocol, as a measure of habituation. Vehicle-treated 4cG mice displayed an absence of startle habituation compared to WT controls, indicating a reduced adaptive response to repeated stimuli **(Fig. 2C)**. In contrast, 4cG mice treated with CP2 for 8 weeks exhibited a restoration of the startle habituation effect **(Fig. 2C)**.

**Figure 2.**
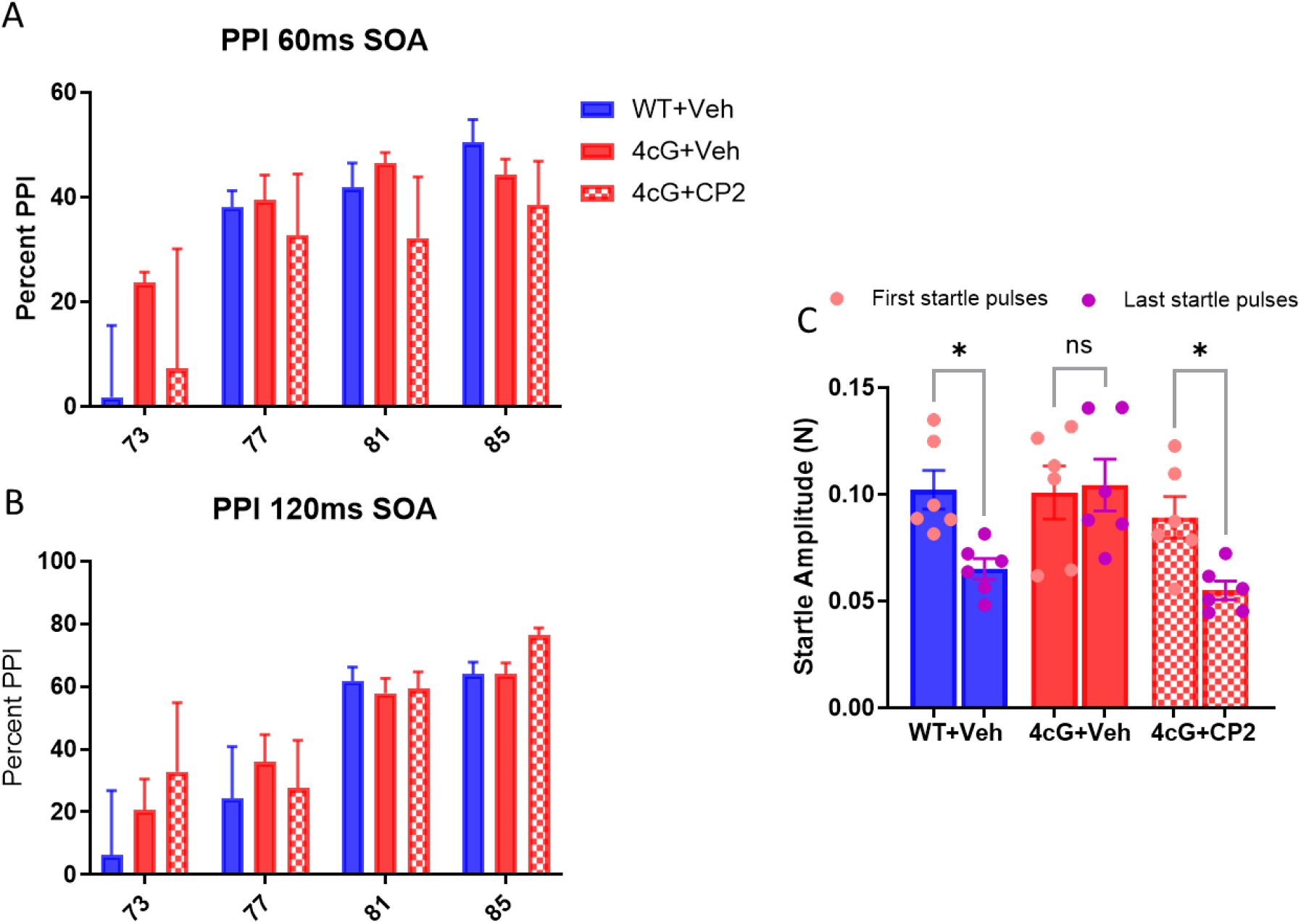
Effect of chronic CP2 treatment on the prepulse inhibition and startle habituation in 4cG mice. **A, B.** The CP2 treatment did not alter prepulse inhibition to acoustic startle (PPI) responses at different prepulse intensities with stimulus onset asynchrony (SOA) of 60 ms and 120 ms in WT mice and in 4cG mice administered vehicle or CP2 for 8 weeks. **C**. Assessment of start habituation. The response to the startle pulses presented before and after the prepulse inhibition experiment showed that vehicle-treated 4cG mice did not show a habituation effect, but the vehicle-treated WT mice and CP2-treated 4cG mice displayed habituation to the startle stimuli. A one-way ANOVA (F (5,32): 5.713, p=0.0008) followed by Bonferroni’s multiple comparisons test revealed a significantly decreased startle response to the last startle pulses compared to the first startle pulses, indicating the habituation effect in vehicle-treated WT mice (t= 2.816, p= 0.0253) and in CP2-treated 4cG mice (t= 2.597, p= 0.0427). WT+Veh: n=5; 4cG+Veh: n=9, 4cG+CP2; n=5. All data presented as mean ± SEM.

We next examined the effect of treatment on social interaction behavior using a three-chamber social interaction paradigm. Vehicle-treated 4cG mice showed reduced preference for social interaction, consistent with previously reported social deficits in 4cG mice^13^. CP2 treatment selectively improved social behavior in 4cG mice. The sociability deficit was not rescued by the CP2 treatment in 4cG mice (**Fig. 3A**). However, in the social novelty test, CP2 (25 mg/kg, 8 weeks) significantly improved the preference for the novel stimulus (S2) compared to the familiar stimulus (S1) in 4cG mice (**Fig. 3B**).

**Figure 3.**
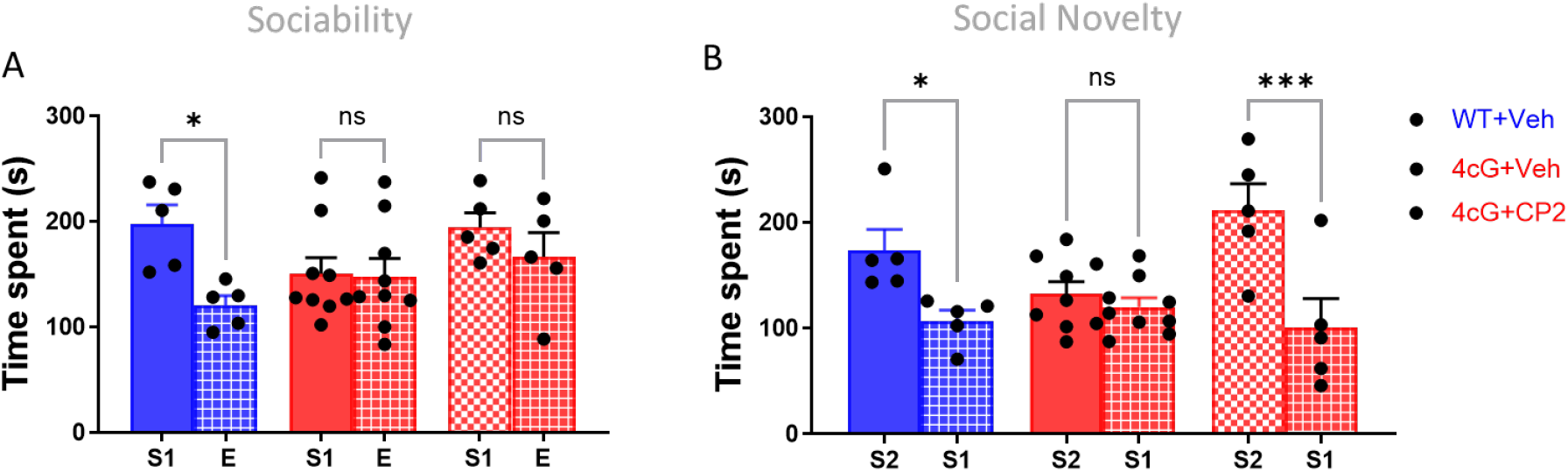
Chronic CP2 treatment improves the behavior of 4cG mice in the social novelty test. **A**. The sociability test was performed with vehicle-treated WT, and vehicle- or CP2-treated 4cG mice. **A** one-way ANOVA (F (5,32): 3.369, p=0.0166) followed by Bonferroni’s multiple comparisons test revealed an increased exploration of a stranger **(S1)** compared to the object **(E)** in vehicle-treated WT mice (t= 3.003, p= 0.0166) but not in vehicle- or CP2-treated 4cG mice. **B**. Subsequently, the vehicle-treated WT, and the vehicle- or CP2-treated 4cG mice were subjected to social novelty preference investigation. A one-way ANOVA (F (5,32): 5.937, p=0.0005) followed by Bonferroni’s multiple comparisons test revealed an increased exploration of stranger **S2** compared to now familiar **S1** in vehicle-treated WT mice (t= 2.573, p= 0.0452) and in CP2-treated 4cG mice (t= 4.272, p= 0.0004). WT+Veh: n=5; 4cG+Veh: n=9, 4cG+CP2: n=5. All data presented as mean ± SEM.

We have previously demonstrated that acute gavage with CP2 induced multiple neuroprotective mechanisms in the hippocampal tissue of APP/PS1 mice via AMPK activation^22^. Upregulation of the expression of the master regulator of mitochondrial biogenesis, PGC-1α^20^, is AMPK-dependent. To investigate whether acute administration of CP2 influences biochemical pathways regulating mitochondrial biogenesis and function in 4cG mice, we administered a single oral dose of CP2 (25 mg/kg) and evaluated protein expression levels 24 hours later. Western blot analyses revealed a significant reduction in levels of PGC-1α in 4cG mice **(Fig. 4A**). Consistent with the mechanism of action, acute CP2 treatment restored the expression of PGC-1α in 4cG mice (**Fig. 4A**). Treatment also resulted in improved expression of VDAC1 in 4cG mice **(Fig. 4A, 4B)**.

**Figure 4.**
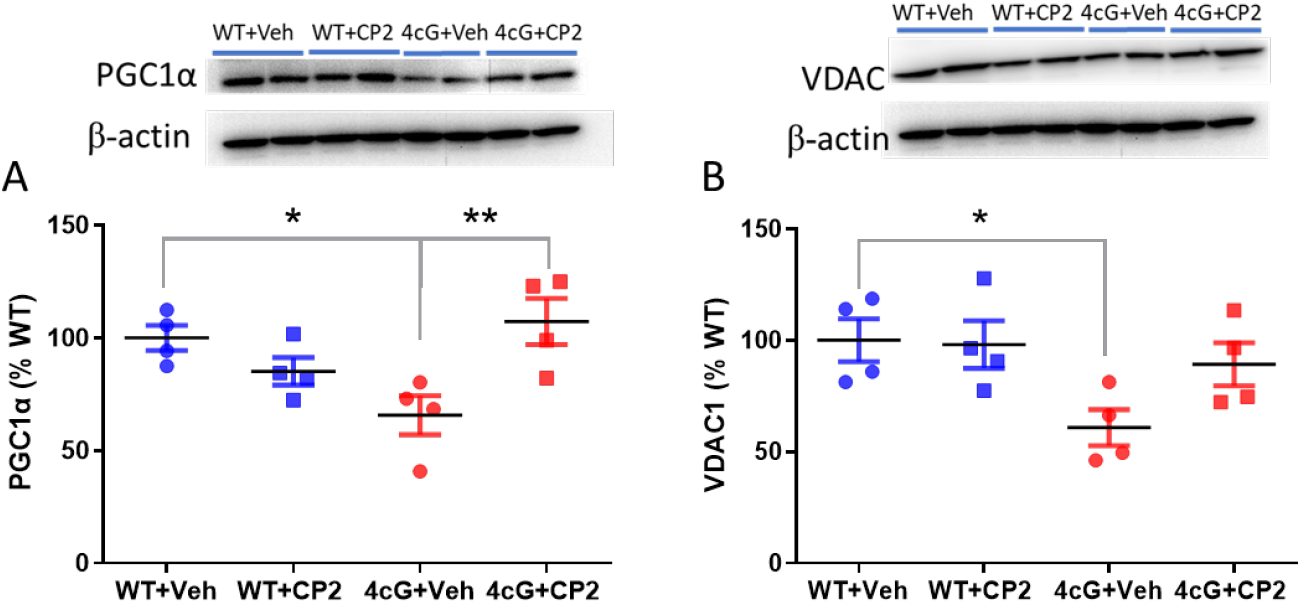
Acute CP2 treatment restores hippocampal protein expression after 24 hours. **A.** Acute CP2 treatment via oral gavage (single dose of 25 mg/kg) rescued PGC-1a protein expression levels in the hippocampus, after 24 hours in 4cG mice. One-way ANOVA (F(3,12): 5.468, p=0.0133) followed by Bonferroni’s multiple comparisons test revealed a difference in PGC-1a expression between vehicle-treated 4cG mice and vehicle-treated WT mice (t= 3.090, p= 0.0281) and an improvement in PGC-1a expression in CP2-treated 4cG mice compared to vehicle-treated 4cG mice to (t=3.744, P=0.0084). **B**. Acute CP2 treatment showed a trend towards increased VDAC1 levels in 4G mice. A one-way ANOVA (F(3,12): 3.572, p=0.0471) followed by Bonferroni’s multiple comparisons test revealed a significant difference in VDAC1 expression between vehicle-treated 4cG mice and vehicle-treated WT mice (t= 2.894, p= 0.0399). WT+Veh: n=4; 4cG+Veh: n=4, 4cG+CP2: n=4. All data presented as mean ± SEM.

Finally, we investigated whether reversals of behavioral deficits were associated with changes in pathways regulating mitochondrial biogenesis. 4cG mice displayed reduced expression of the key regulators of mitochondrial biogenesis and function, PGC-1α and VDAC1, in the hippocampus compared to WT controls (**Fig. 5**). Chronic CP2 treatment (25 mg/kg, 8 weeks) significantly increased the expression of these key mitochondrial markers in the hippocampus of 4cG mice. Specifically, levels of the master regulator of mitochondrial biogenesis, PGC-1α, and of the mitochondrial outer membrane protein VDAC1 were both reached levels comparable to those of wild-type controls (**Fig. 5A, 5B**). These results suggest that CP2 enhances mitochondrial function and biogenesis, potentially underlying the observed behavioral rescue.

**Figure 5.**
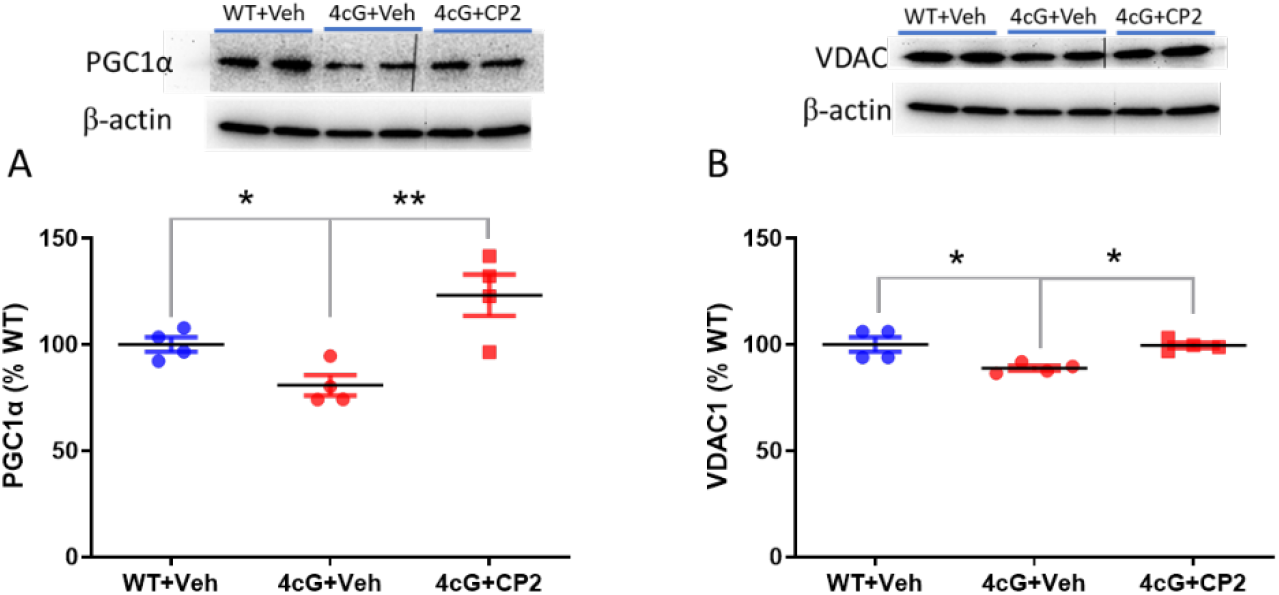
Chronic CP2 treatment restores the expression of PGC-1a and VDAC1 in the hippocampus of 4cG mice. **A.** Chronic CP2 treatment restored the protein expression of the master regulator of mitochondrial biogenesis PGC-1a in 4cG mice. A one-way ANOVA (F(2,9): 10.44, p=0.0045) followed by Bonferroni’s multiple comparisons test revealed a significant difference in PGC-1a expression between vehicle-treated 4cG mice and vehicle-treated WT mice (t= 3.090, p= 0.0281) and PGC-1a expression was increased in CP2-treated 4cG mice compared to vehicle-treated 4cG mice (t=3.744, P=0.0084). **B**. The mitochondrial outer membrane protein voltage-dependent anion channel 1 (VDAC1) expression was also increased with chronic CP2 treatment in 4cG mice. A one-way ANOVA (F(2,9): 7.848, p=0.0106) followed by Bonferroni’s multiple comparisons test revealed a significant reduction in VDAC1 expression in vehicle-treated 4cG mice compared to vehicle-treated WT mice (t= 3.487, p= 0.0206) and an improvement in VDAC1 expression in CP2-treated 4cG mice compared to vehicle-treated 4cG mice (t=3.373, P=0.0247). WT+Veh: n=4; 4cG+Veh: n=4, 4cG+ CP2, n=4. All data presented as mean ± SEM.

Together, these data are consistent with CP2 activating AMPK and downstream mitochondrial biogenesis pathways under both acute and chronic (8-week) treatment conditions. This sustained activation is associated with improved cognitive function and a rescue of behavioral deficits in 4cG mice.

## Discussion

The present study demonstrates that chronic administration of CP2 for 8 weeks produces a robust behavioral and molecular rescue of deficits in 4cG mice, a model for schizophrenia characterized by cognitive, sensorimotor, and social deficits. Specifically, CP2 restored working memory, normalized startle habituation, improved social preference, and selectively enhanced the expression of key proteins of mitochondrial regulatory pathways in the hippocampus. These findings align with a growing body of evidence implicating mitochondrial dysfunction as an important driver of neurodevelopmental phenotypes and CP2 as a potential therapeutic compound for schizophrenia.

CP2 is a brain-penetrant, weak mitochondrial complex I inhibitor that induces adaptive bioenergetic responses rather than overt mitochondrial suppression^21^. Weak inhibition of complex I has been shown to increase the AMP/ATP ratio, activating AMPK-dependent pathways that promote neuroprotection and metabolic adaptation^20,24^. Consistent with this mechanism, our findings of improved behavioral outcomes following chronic CP2 treatment suggest that metabolic reprogramming can effectively reverse functional deficits in 4cG mice.

A key observation in this study is the upregulation of PGC-1α in the hippocampus following CP2 treatment. PGC-1α is widely recognized as a master regulator of mitochondrial biogenesis and cellular energy metabolism^33,34^. Its activation has been linked to improved neuronal resilience, enhanced oxidative phosphorylation, and mitochondrial biogenesis. Previous studies have shown that increasing PGC-1α expression mitigates mitochondrial dysfunction and improves cognitive outcomes in models of brain injury and neurodegenerative disease^19–21^. Therefore, the observed restoration of PGC-1α expression in both acute and chronic CP2-treated 4cG mice provides a mechanistic link between CP2 and mitochondrial remodeling. Although both acute and chronic treatments produced biochemical restoration, only chronic, sustained treatment was sufficient to translate these changes into behavioral rescue.

In parallel, CP2 treatment increased VDAC1 expression, a key component of the mitochondrial outer membrane that regulates metabolite exchange and cellular energy homeostasis. VDAC1 has a complex role in brain pathology; its dysregulation has been implicated in diseases such as Alzheimer’s disease^35^ and schizophrenia^36^. While excessive or pathological VDAC1 activity may contribute to mitochondrial toxicity, its proper expression is essential for maintaining mitochondrial function and metabolic flux. The increase in VDAC1 observed here likely reflects improved mitochondrial integrity and metabolic capacity, rather than pathological overactivation, particularly in the context of concurrent PGC-1α upregulation.

Behaviorally, the rescue of working memory, sensory habituation, and social interaction suggests that mitochondrial dysfunction contributes broadly to neural circuit impairments in 4cG mice.

These findings are consistent with literature demonstrating that mitochondrial deficits disrupt synaptic plasticity, neurotransmission, and network-level activity, ultimately leading to cognitive and behavioral abnormalities^13,17,37,38^. Importantly, the progressive improvement observed over 8 weeks indicates that sustained metabolic intervention is required to achieve functional recovery, emphasizing the importance of chronic administration.

This study provides proof-of-concept evidence for CP2 effects in a mouse model for schizophrenia, but several limitations should be acknowledged. Future studies could investigate downstream signaling pathways linking CP2-dependent modulation of mtCI and PGC-1α activation, including AMPK and NAD^+^-dependent pathways. Assessing mitochondrial dynamics, respiration, and synaptic function will further strengthen our knowledge of the cellular basis of behavioral rescue. While 8 weeks of treatment were effective, it remains unclear whether alternative dosing regimens could yield greater or faster benefits. Longitudinal studies examining treatment initiation at different developmental (life) stages will be particularly informative.

## Conclusions

In summary, this study identifies chronic CP2 treatment as an effective strategy to reverse behavioral and mitochondrial deficits in 4cG mice. By enhancing PGC-1α–mediated mitochondrial biogenesis and restoring VDAC1 expression, CP2 promotes a bioenergetic state that supports neuronal function and behavioral performance. Suggesting that the selective targeting of mitochondrial pathways is sufficient to achieve therapeutic benefit.

These findings are significant because they provide direct evidence that modulating mitochondrial function can reverse complex behavioral phenotypes, rather than merely slowing disease progression. Given the limited availability of disease-modifying therapies for disorders associated with mitochondrial dysfunction, CP2 represents a promising candidate for translational development. CP2 has already demonstrated efficacy in preclinical models of neurodegeneration, including Alzheimer’s and Huntington’s disease^25,39^. Extending these findings to other neurodevelopmental and metabolic disorders will help define the broader applicability of this approach.

The ability of CP2 to cross the blood–brain barrier and to selectively modulate mitochondrial function positions this compound, or optimized next-generation compounds such as C458^21^ and C273^40^, as a strong candidate for clinical development. Targeting mitochondrial bioenergetics represents a unifying therapeutic strategy for disorders characterized by impaired neuronal metabolism, including schizophrenia and neurodegenerative diseases, and potentially also autism spectrum disorders.

## Acknowledgements

This study was supported by institutional startup funds provided by the College of Veterinary Medicine of the University of Illinois Urbana-Champaign (to UR) and by grant R01AG055549 from the National Institute of Health (to ET).

## Data availability

Available upon request.

